# Oncogenic NPM-ALK reprograms the TGM1 interactome toward oncogenic signaling and transcriptional states

**DOI:** 10.64898/2026.08.01.742203

**Authors:** So Taguchi, Kosuke Higashi, Yuuki Tanaka, Hidetaka Kosako, Kazumasa Aoyama

## Abstract

Oncogenic NPM-ALK drives aberrant signaling networks that promote malignant phenotypes; however, the molecular mechanisms linking oncogenic signaling to downstream cellular programs remain incompletely understood. Among candidate regulatory factors, transglutaminase 1 (TGM1) has not been functionally characterized in this context. Here, we investigated the role of TGM1 in NPM-ALK–expressing cells by combining proximity-dependent proteomics with functional analyses. Using a TurboID-based approach, we mapped the TGM1-associated protein network and identified extensive remodeling of this network upon NPM-ALK expression. Proteomic analyses revealed that NPM-ALK reduced TGM1-associated proteins involved in genome maintenance and DNA repair, while enhancing associations with proteins linked to cytoplasmic translation and PI3K–AKT signaling pathways. Consistent with these findings, TGM1 deficiency impaired cell proliferation without significantly affecting cell viability, indicating a specific role in maximal proliferative capacity. Furthermore, proteomic and functional analyses suggested a link between TGM1 and AKT signaling pathways. Together, these findings suggest that oncogenic NPM-ALK reprograms the TGM1 interactome toward oncogenic signaling and transcriptional states, positioning TGM1 within signaling networks associated with proliferative cellular phenotypes.

## Introduction

The NPM-ALK fusion protein, generated by the t(2;5)(p23;q35) chromosomal translocation, is a constitutively active tyrosine kinase that drives oncogenesis in anaplastic large cell lymphoma (ALCL) [1]. NPM-ALK activates multiple downstream signaling pathways, including STAT3, PI3K–AKT, and MAPK signaling, thereby promoting malignant proliferation, survival, and transcriptional reprogramming [2–4]. Although the downstream signaling mechanisms of NPM-ALK have been extensively studied, how oncogenic signaling reshapes protein interaction networks linked to cellular phenotypes remains incompletely understood.

Protein interaction networks play central roles in organizing intracellular signaling and determining cellular responses to oncogenic stimuli. Recent advances in proximity-dependent labeling approaches, including TurboID-based proteomics, have enabled comprehensive characterization of dynamic protein-associated networks in living cells [5, 6]. Indeed, recent studies have demonstrated that oncogenic NPM-ALK can selectively remodel protein interaction networks, including the MRVI1 interactome [7]. Among candidate regulatory molecules, transglutaminase family proteins are multifunctional enzymes involved not only in protein crosslinking but also in the regulation of cellular signaling and structural organization [8, 9]. TGM1, a member of the transglutaminase family, is best characterized for its role in epidermal differentiation and cornified envelope formation [10–13]. In contrast to TGM2, which has been implicated in tumor progression and cancer cell survival [14–16], the functional roles of TGM1 in oncogenic signaling and proliferative regulation remain poorly understood, despite recent studies suggesting associations with human malignancies [17].

In this study, we investigated the role of TGM1 in NPM-ALK–expressing cells by combining proximity-dependent proteomics with functional analyses. Using a TurboID-based approach, we characterized TGM1-associated protein networks and examined how these networks are remodeled by oncogenic NPM-ALK signaling. In addition, we evaluated the effects of TGM1 deficiency on AKT signaling, proliferative capacity, and transcriptional programs. Our findings suggest that oncogenic NPM-ALK reprograms the TGM1 interactome toward oncogenic signaling and transcriptional states, positioning TGM1 within signaling networks required for maximal proliferative capacity.

## Materials and Methods

### Cells and transfection

Lenti-X 293T cells (#632180, Clontech) [7] were cultured in DMEM (high glucose, 4.5g/L Glucose) supplemented with 10% fetal bovine serum (FBS), 100 U/mL penicillin, and 100 μg/mL streptomycin. Ba/F3 cells and Ba/F3 cells expressing NPM-ALK [18] were cultured in DMEM (high glucose, 4.5g/L Glucose) supplemented with 10% fetal bovine serum (FBS), 100 U/mL penicillin, 100 μg/mL streptomycin, and 2 ng/mL recombinant murine IL-3. Cells were maintained at 37°C in a humidified incubator with 5% CO₂. Transfection was conducted using polyethyleneimine.

### Plasmids

The CSII-V5-TurboID-TGM1-IRES-Puro vector was generated by inserting the TGM1 cDNA obtained by reverse transcription and PCR amplification from mRNA of NPM-ALK-expressing Ba/F3 cells [18] into the CSII-V5-TurboID-IRES-Puro vector [7, 19]. The PCR was performed using the following primers: forward, 5’-gatgagtttgaatatgatgagctgattgtgcgccg-3’ and reverse, 5’-gtttggaacttccacgtgtggaacgactgctggat-3’. The MSCV-NPM-ALK-IRES-GFP vector previously constructed [7, 18] was used in this study.

TGM1 knockout was performed using the CRISPR-Cas9 system as described previously [20] with minor modifications. Single guide RNA (sgRNA) sequences were designed using the CHOPCHOP web tool (https://chopchop.cbu.uib.no/) and cloned into the lentiCRISPR v2 vector (Addgene, Plasmid #52961). Lentiviral particles were produced in LentiX 293T cells and used to infect NPM-ALK–expressing Ba/F3 cells to introduce sgRNAs. Following puromycin selection, TGM1 knockout cells were established. The sgRNA sequences used in this study were as follows:

5’-GATCCACTCCAACAATCGAG-3’ (sgTGM1#1)
5’-TTCGAGCCGGAAGACGACGT-3’ (sgTGM1#2)
5’-CAGCTTTAAGATAGTGTACG-3’ (sgTGM1#3)

### Antibodies

The following antibodies were used: anti-TGM1 (#12912-3-AP, Proteintech), anti-Phospho-AKT (Ser473) (#4060, Cell Signaling Technology), anti-AKT (#sc-5298, Santa Cruz Biotechnology) anti-V5 (#M215-3, Medical & Biological Laboratories; #sc-271944, Santa Cruz Biotechnology), anti-ALK (#3633, Cell Signaling Technology) anti-Tubulin (#MCA77G, Bio-Rad), anti-Actin (#sc47778, Santa Cruz Biotechnology; #4970, Cell Signaling Technology), Anti-Mouse IgG-HRP (#NA931, GE Healthcare; light chain specific, #115-035-174, Jackson), Anti-Rabbit IgG-HRP (#NA934, GE Healthcare; light chain specific, #211-032-171, Jackson), Anti-Rat IgG-HRP (#sc-2006, Santa Cruz; #7077, Cell Signaling Technology), and Normal Rabbit IgG (#2729, Cell Signaling Technology) antibodies.

### Biotin-based proximity labeling proteomic analysis

Proximity-dependent biotin labeling combined with mass spectrometry was performed to characterize proteins associated with TGM1, as described previously with minor modifications [7]. LentiX-293T cells were transiently transfected with V5-TurboID or V5-TurboID-TGM1 in the presence or absence of NPM-ALK. To induce proximity-dependent biotinylation, cells were incubated with 500 μM D-biotin for the final 24 h of a 48 h culture period. Cells were harvested and lysed in guanidine-containing buffer (6 M guanidine-HCl, 100 mM HEPES-NaOH, pH 7.5) supplemented with 10 mM tris(2-carboxyethyl)phosphine (TCEP) and 40 mM chloroacetamide (CAA). Biotinylated proteins were affinity-purified using Tamavidin 2-REV beads [21, 22] and analyzed by liquid chromatography–tandem mass spectrometry (LC-MS/MS). Proteomic data were processed using Proteome Discoverer software (Thermo Fisher Scientific). Proteins enriched in V5-TurboID-TGM1 samples relative to the TurboID control were defined based on abundance ratios and statistical significance thresholds.

### Western blotting

Western blot analysis was carried out essentially as described previously [23, 24]. Cells were washed twice with PBS and lysed in SDS-containing lysis buffer (2% SDS, 20 mM Tris-HCl, pH 8.0) [25]. Cell lysates were sonicated, mixed with an equal volume of 2× SDS sample buffer, and heated at 95°C for 10 min. Proteins were separated by SDS-PAGE and transferred onto PVDF membranes (PALL). After blocking with 1% BSA, membranes were incubated with the indicated primary and secondary antibodies. Biotinylated proteins were detected using HRP-conjugated streptavidin (#405210, BioLegend). Signals were visualized using Immobilon Western chemiluminescence reagent (Millipore) and acquired with a ChemiDoc™ Touch imaging system (Bio-Rad). For sequential reprobing, antibodies were removed by incubation in 0.2 M glycine-HCl buffer (pH 2.5), and/or HRP activity was quenched using 0.1% NaN3, as described previously [23, 26]. Image processing for figure preparation was performed using GIMP software, and densitometric analysis of band intensities was conducted using ImageJ software [27].

### Immunofluorescence microscopy

Immunofluorescence analysis was carried out largely according to previously described procedures [28, 29] with minor modifications. Cells were seeded onto poly-L-lysine– coated glass coverslips (22 × 22 mm, Matsunami) and transfected with the indicated plasmids. Following incubation with 500 µM biotin for 24 h, cells were fixed with 4% paraformaldehyde for 20 min and washed three times with PBS. Cells were subsequently permeabilized using 0.5% Triton X-100 in PBS containing 1% BSA for 20 min. The coverslips were incubated overnight at 4°C with anti-V5 antibody diluted in blocking buffer. Biotinylated proteins were simultaneously detected using CoraLite® Plus 488-conjugated streptavidin (#PF00023, Proteintech). After washing with PBS, Multi-rAb™ CoraLite® Plus 594-Goat Anti-Rabbit Recombinant Secondary Antibody (H+L) (#RGAR004, Proteintech) was applied for 30–60 min at room temperature in the dark. Samples were mounted using Fluoro-KEEPER Antifade Reagent, Non-Hardening Type with DAPI (Nacalai Tesque), and fluorescence images were acquired using an FV4000 confocal laser scanning microscope (Evident, Japan).

Quantification of fluorescence signal intensities was performed using ImageJ software (NIH), essentially as described previously [30, 31]. Nuclear and whole-cell regions were manually defined based on DAPI staining and cell morphology, respectively. Mean fluorescence intensities of V5 and biotin signals were measured for each individual cell, and nuclear-to-whole cell intensity ratios were calculated.

### RNA-seq analysis

Total RNA sequencing was performed by Kazusa DNA Research Institute (Chiba, Japan). Ribosomal RNA was depleted using the NEBNext rRNA Depletion Kit v2 (Human/Mouse/Rat) (New England Biolabs), and sequencing libraries were prepared using the NEBNext Ultra II Directional RNA Library Prep Kit for Illumina according to the manufacturer’s protocols. Sequencing was carried out on an Illumina NextSeq 500 platform using single-read 75 bp conditions. Sequence reads were mapped to the mouse genome (mm10) using HISAT2 v2.2.1, and FPKM values were calculated using Cufflinks v2.2.1 by Kazusa DNA Research Institute.

### Gene Ontology (GO) Enrichment Analysis

Functional enrichment analysis was conducted using the Database for Annotation, Visualization, and Integrated Discovery (DAVID) platform [32, 33], as described previously [7, 34] with minor modifications. Lists of proteins or differentially expressed genes were analyzed using official gene symbols, and enrichment was evaluated within the Gene Ontology (GO) Biological Process category under default human background settings. GO terms with p-values below 0.05 were regarded as significantly enriched. The resulting annotation tables were exported and processed using Microsoft Excel. Enriched GO categories were subsequently visualized as bar graphs.

### Gene set enrichment analysis (GSEA)

Gene set enrichment analysis (GSEA) was performed using the Broad Institute GSEA software, as described previously [35, 36]. Differentially expressed genes identified from RNA-seq analysis were ranked according to expression changes between control and TGM1-knockout cells. Hallmark gene sets from the Molecular Signatures Database (MSigDB) were used for pathway enrichment analysis[37].

### Structural analysis

Structural visualization and annotation of human AKT1 were performed using *The PyMOL Molecular Graphics System* (Schrödinger, LLC) based on the crystal structure deposited in the Protein Data Bank (PDB ID: 6NPZ). The analyzed structure contains the protein kinase domain and C-terminal regulatory region of human AKT1. AKT1 amino acid sequence information was obtained from UniProt (P31749-1). AKT1-derived peptides identified by proximity labeling analysis were mapped onto the primary amino acid sequence of AKT1. The protein kinase domain, C-terminal tail, Ser473 residue, and the putative TGM1 interaction region were annotated and visualized on the structure. The spatial relationship between the kinase domain-associated region and the Ser473-containing C-terminal tail was assessed using the displayed structure.

### Cell proliferation and viability assays

Cell proliferation assays were performed using Ba/F3 cells expressing NPM-ALK, as described previously with minor modifications [7, 38]. Cells were seeded at equal densities and cultured for the indicated periods. Cell numbers were measured at each time point using a hemocytometer, and proliferation rates were calculated relative to day 0 or to control NPM-ALK–expressing cells, as indicated in each figure. For viability assays, cells were cultured for 24–72 h, stained with trypan blue solution, and viable cells were quantified manually using a hemocytometer.

### Deposition of data

The mass spectrometry data have been deposited to jPOST (JPST003926) and ProteomeXchange (PXD065924). RNA-seq data have been deposited in the NCBI Sequence Read Archive (SRA) under accession number PRJNA1464642.

## Results

### Establishment of a proximity labeling system to characterize the TGM1 interactome

To systematically characterize the molecular environment surrounding TGM1, we established a proximity-dependent biotinylation system using a V5-tagged TurboID fused to TGM1. Expression of V5-TurboID-TGM1 in LentiX cells resulted in robust biotinylation of proximal proteins in a time-dependent manner upon biotin supplementation, whereas minimal labeling was observed in cells expressing TurboID alone (Fig. 1A, B). Immunofluorescence analysis revealed that the subcellular distribution of TurboID-TGM1 was consistent with previously reported localization patterns of endogenous TGM1, as documented in public databases, including the Human Protein Atlas[39], supporting the physiological relevance of the fusion construct. In contrast, TurboID alone exhibited a diffuse distribution, confirming that proximity labeling was dependent on TGM1 localization (Fig. 1C, D). Together, these results validate the TurboID-based approach as a robust platform for profiling the TGM1-associated protein network and provide the basis for subsequent interactome analyses.

**Figure 1.**
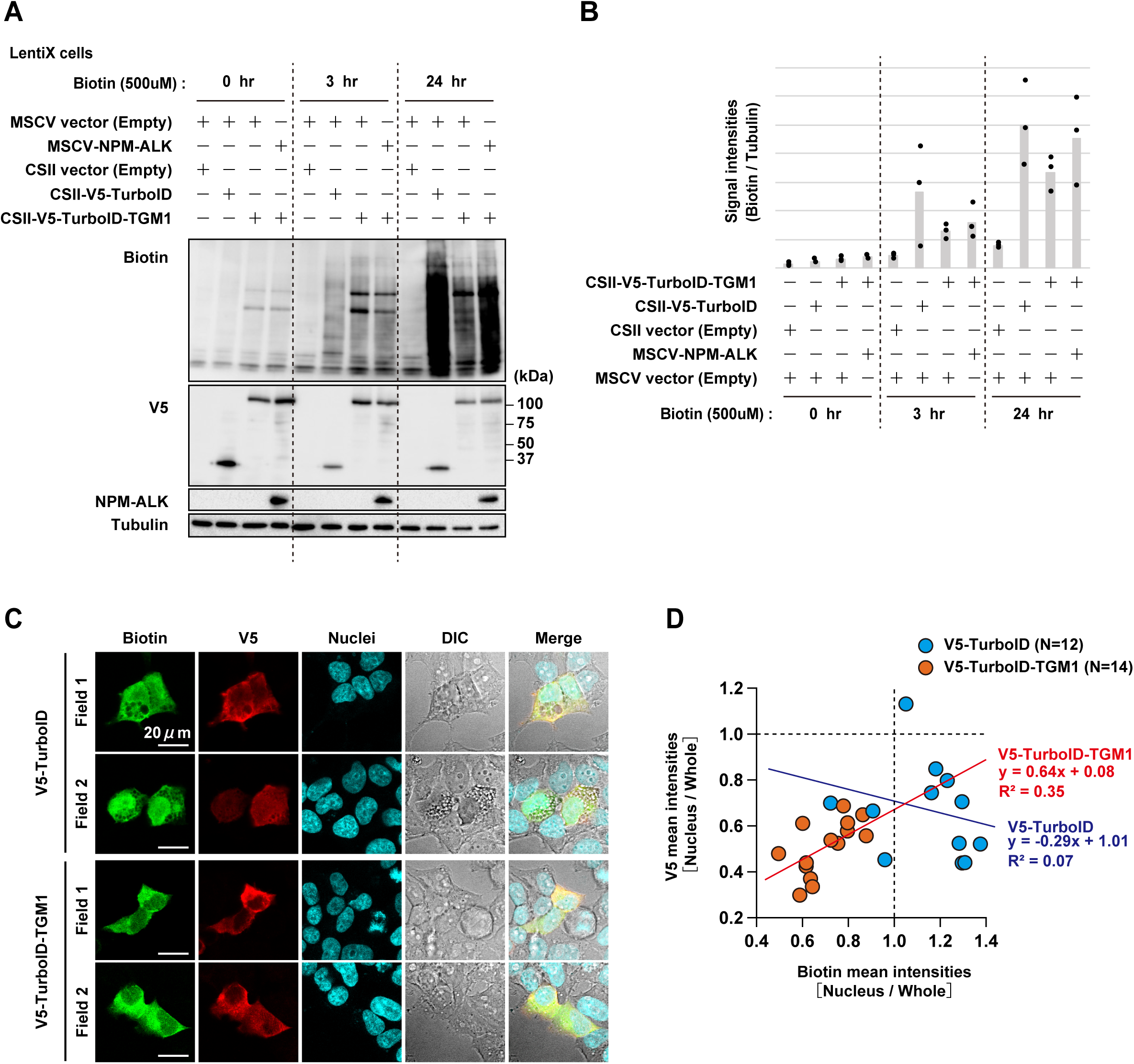
Validation of TurboID-TGM1 proximity labeling system. (A) Western blot analysis showing biotinylation of proteins in LentiX cells transiently transfected with V5-TurboID or V5-TurboID-TGM1 together with MSCV vector (empty) or MSCV-NPM-ALK. Cells were incubated with biotin for the indicated times. Tubulin was used as a loading control. (B) Quantification of biotinylation signals normalized to tubulin. Data represent relative signal intensities at the indicated time points. (C) Immunofluorescence images of LentiX cells expressing V5-TurboID or V5-TurboID-TGM1. Cells were stained for V5 and biotinylated proteins. Representative images are shown. Scale bars, 20 μm. (D) Quantification of nuclear-to-whole cell intensity ratios for V5 and biotin signals. Each dot represents an individual cell.

### Global mapping of the TGM1 interactome reveals a cytoskeleton-associated protein network

To define the molecular network associated with TGM1, we performed proximity labeling–based proteomic analysis using TurboID-TGM1. Across independent replicates, a total of 1,590 proteins were reproducibly identified as TGM1-associated candidates based on enrichment over TurboID control (Fig. 2A-C). Gene ontology analysis revealed that these proteins were significantly enriched in pathways related to cytoskeleton organization, intracellular signaling, and protein modification processes (Fig. 2D). Consistent with this finding, multiple cytoskeleton-associated factors, including ARPC5, MAP7D2, and ANK2, were enriched in the TGM1-associated network (Fig. 2E), supporting a role for TGM1 in structural and spatial organization within the cell. These results indicate that TGM1 is embedded in a broad protein network linked to cytoskeletal architecture and intracellular signaling, providing a structural framework for subsequent analysis of oncogenic reprogramming. TGM1 is a transglutaminase family protein known to mediate protein crosslinking and structural stabilization during epidermal differentiation and cornified envelope formation [10–13]. These findings are consistent with the established roles of TGM1 in structural organization and protein crosslinking, while additionally suggesting broader functions in intracellular signaling-associated protein networks.

**Figure 2.**
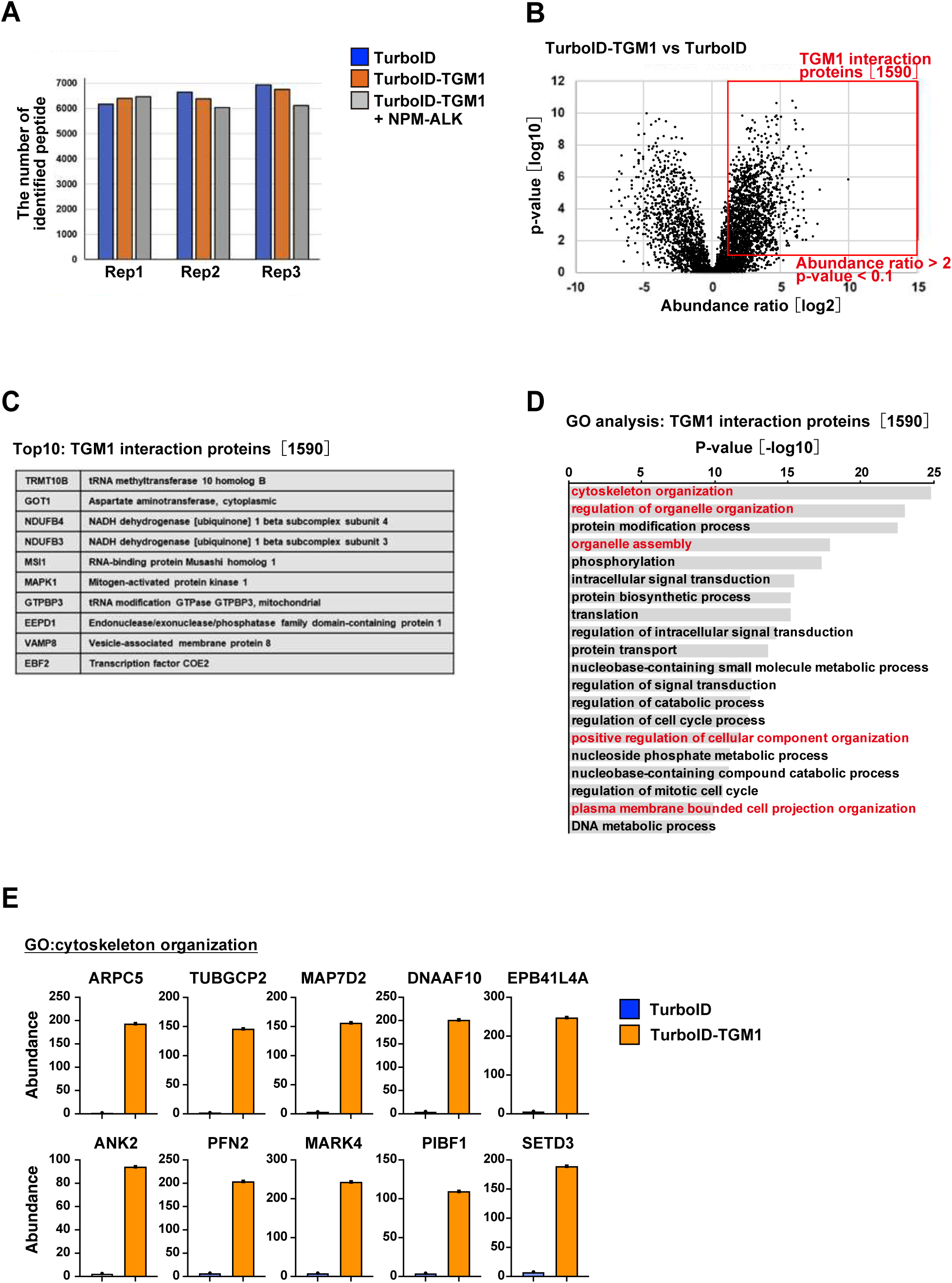
Identification of TGM1-associated proteins. (A) Number of identified peptides in proximity labeling experiments using TurboID or TurboID-TGM1 in the presence or absence of NPM-ALK across independent replicates. (B) Scatter plot showing protein abundance in TurboID-TGM1 versus TurboID samples. Proteins enriched in the TurboID-TGM1 condition were defined based on abundance ratio (>2) and statistical significance (p < 0.1). (C) Top enriched TGM1-associated proteins identified by proximity labeling. Representative proteins are shown. (D) Gene ontology (GO) analysis of TGM1-associated proteins, highlighting enrichment in cytoskeletal organization, intracellular signaling, and protein modification pathways. (E) Representative examples of cytoskeleton-related proteins enriched in the TGM1-associated network. Protein abundance is shown for TurboID and TurboID-TGM1 conditions.

### Oncogenic NPM-ALK suppresses tumor-suppressive components of the TGM1 interactome

To determine how oncogenic signaling reshapes the TGM1-associated protein network, we next examined alterations in the TGM1 interactome upon NPM-ALK expression. Among the 1,590 TGM1-associated proteins identified in Fig. 2B, comparative analysis identified a subset of 135 proteins whose association with TGM1 was significantly reduced in the presence of NPM-ALK (Fig. 3A, B). Gene ontology analysis revealed that these downregulated interactors were highly enriched in pathways related to DNA repair, DNA damage response, and p53-associated signaling (Fig. 3C). Representative factors, including REV3L, ERCC2, and GEN1, showed marked decreases in association with TGM1 upon NPM-ALK expression (Fig. 3D). These results indicate that oncogenic NPM-ALK selectively suppresses TGM1 interactions linked to genome maintenance and tumor-suppressive pathways, suggesting a shift in the functional landscape of the TGM1 interactome.

**Figure 3.**
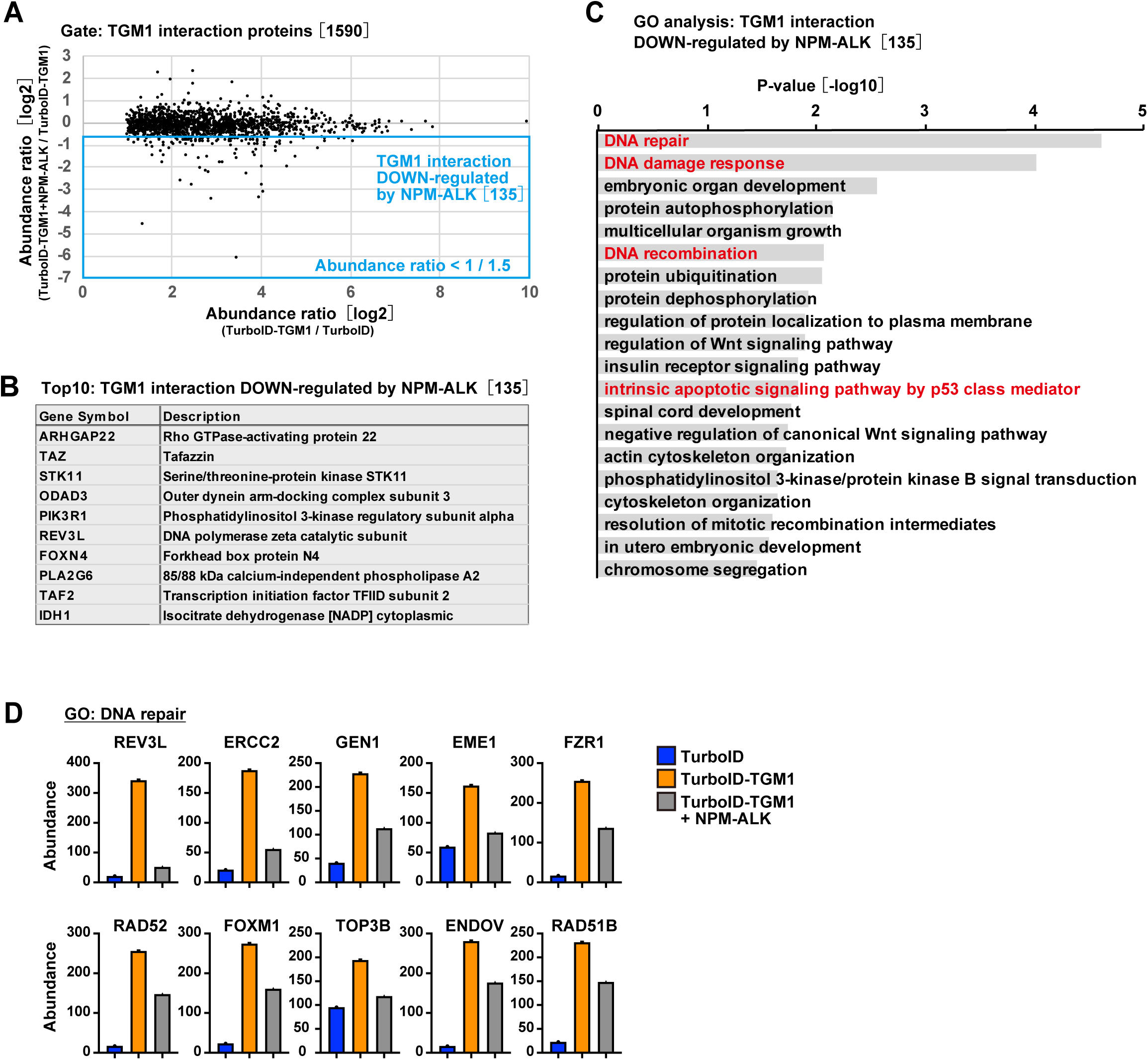
Downregulation of TGM1-associated proteins by NPM-ALK expression. (A) Scatter plot showing changes in protein abundance among TGM1-associated proteins upon NPM-ALK expression. Proteins downregulated by NPM-ALK were defined based on abundance ratio < 1/1.5. (B) Top 10 TGM1-associated proteins downregulated by NPM-ALK expression. (C) Gene ontology (GO) analysis of TGM1-associated proteins downregulated by NPM-ALK expression. Enriched terms related to DNA repair, DNA damage response, DNA recombination, and p53-associated apoptotic signaling are highlighted. (D) Representative DNA repair-related proteins among the TGM1-associated proteins downregulated by NPM-ALK expression. Protein abundance is shown for TurboID, TurboID-TGM1, and TurboID-TGM1 + NPM-ALK conditions.

### Oncogenic NPM-ALK redirects the TGM1 interactome toward oncogenic signaling pathways

In contrast to the loss of tumor-suppressive interactions, we next examined how NPM-ALK expression promotes formation of alternative TGM1-associated protein networks. Among the 1,530 TGM1-associated proteins identified under NPM-ALK–expressing conditions (Fig. 4A), comparative analysis identified a distinct subset of 116 proteins whose association with TGM1 was enhanced upon NPM-ALK expression (Fig. 4B). Gene ontology analysis revealed that these upregulated interactors were significantly enriched in pathways associated with protein translation, phosphorylation, and PI3K– AKT signaling (Fig. 4C, D). Notably, key regulators of growth signaling, including AKT1 and mTOR, exhibited increased association with TGM1 in the presence of NPM-ALK (Fig. 4E). These findings suggest that oncogenic NPM-ALK reprograms the TGM1 interactome by reducing tumor-suppressive interactions while enhancing associations with proteins involved in growth-promoting signaling pathways. Together with the results in Fig. 3, this bidirectional remodeling indicates a coordinated shift of the TGM1-associated network toward an oncogenic state.

**Figure 4.**
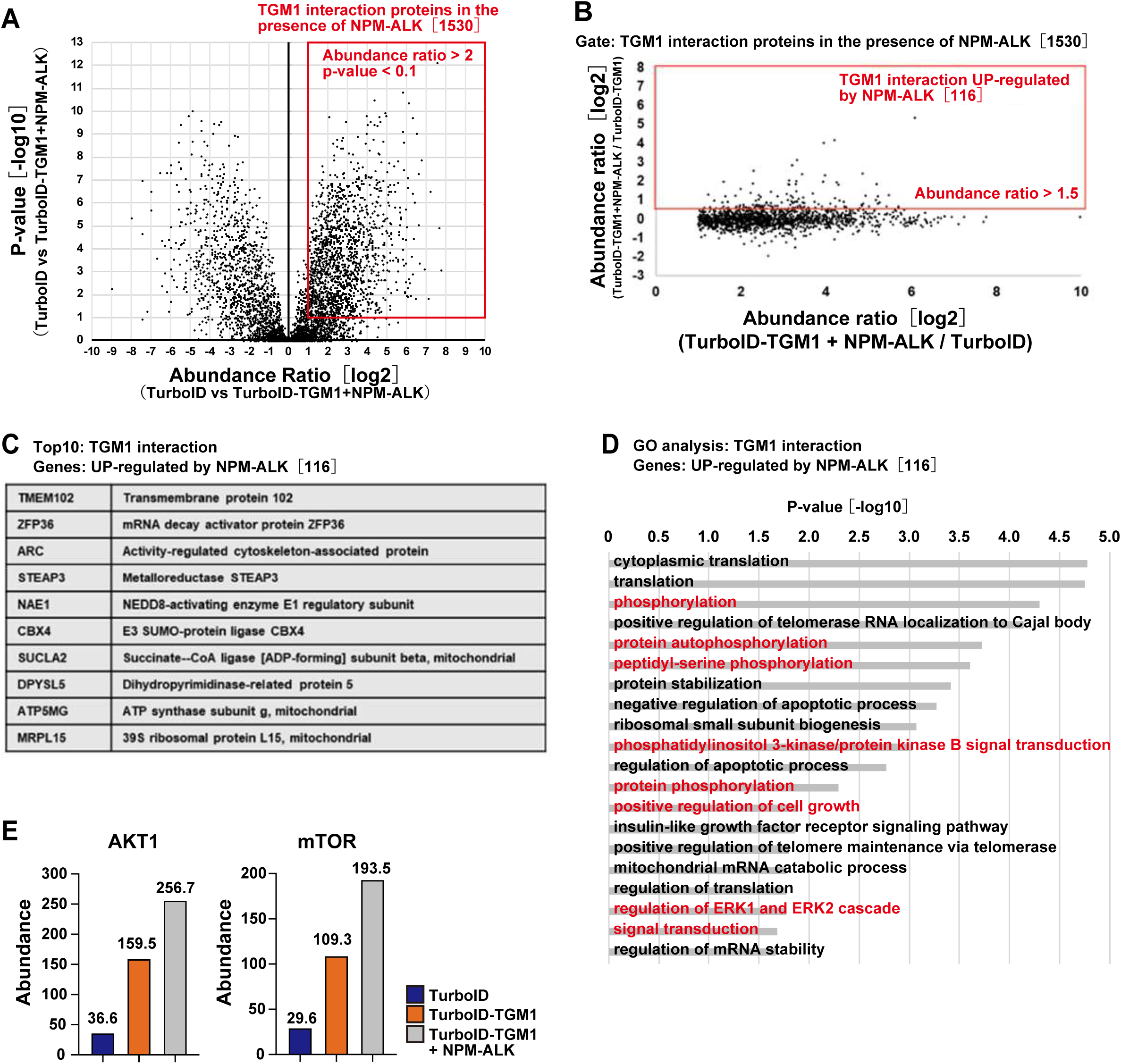
Upregulation of TGM1-associated proteins by NPM-ALK expression. (A) Identification of TGM1-associated proteins in the presence of NPM-ALK. Proteins enriched in TurboID-TGM1 relative to TurboID were defined based on abundance ratio (>2) and statistical significance (p < 0.1), yielding 1,530 TGM1-associated proteins under NPM-ALK–expressing conditions. (B) Scatter plot showing changes in protein abundance within the TGM1-associated proteins identified in (A). Proteins upregulated by NPM-ALK were defined based on abundance ratio (>1.5). (C) Top 10 TGM1-associated proteins upregulated by NPM-ALK expression. (D) Gene ontology (GO) analysis of TGM1-associated proteins upregulated by NPM-ALK, highlighting enrichment in cytoplasmic translation, phosphorylation, and PI3K–AKT signaling pathways. (E) Representative proteins associated with signaling pathways, including AKT1 and mTOR, showing increased abundance in the TGM1-associated network upon NPM-ALK expression.

### TGM1 contributes to AKT signaling in NPM-ALK–expressing cells

We next examined whether the NPM-ALK-dependent enrichment of AKT pathway components in the TGM1 interactome was reflected at the signaling level. Immunoblot analysis showed that TGM1 expression itself was not significantly altered by NPM-ALK expression in Ba/F3 cells (Fig. 5A). Proteomic analysis demonstrated that multiple signaling-associated proteins, including AKT1 and mTOR, were enriched in the TGM1-associated network in the presence of NPM-ALK (Fig. 4D, E). Gene ontology analysis further revealed enrichment of PI3K–AKT signaling-related pathways among proteins whose association with TGM1 was enhanced by NPM-ALK expression (Fig. 4D). Because phosphorylation of AKT at Ser473 is a well-established marker of AKT activation downstream of PI3K signaling [40, 41], these findings raised the possibility that TGM1 contributes to regulation of AKT signaling in NPM-ALK–expressing cells. To functionally investigate this possibility, we generated multiple independent TGM1-knockout Ba/F3 cell clones using a CRISPR-Cas9–based approach, as described in Materials and Methods. Efficient depletion of TGM1 was confirmed in these independent knockout clones (Fig. 5B). Notably, phosphorylation of AKT at Ser473 was reduced in TGM1-deficient cells compared with control NPM-ALK-expressing cells, whereas total AKT levels were not markedly altered (Fig. 5C). These results are consistent with the proteomic data and suggest that TGM1 is linked to AKT signaling in the context of NPM-ALK expression.

**Figure 5.**
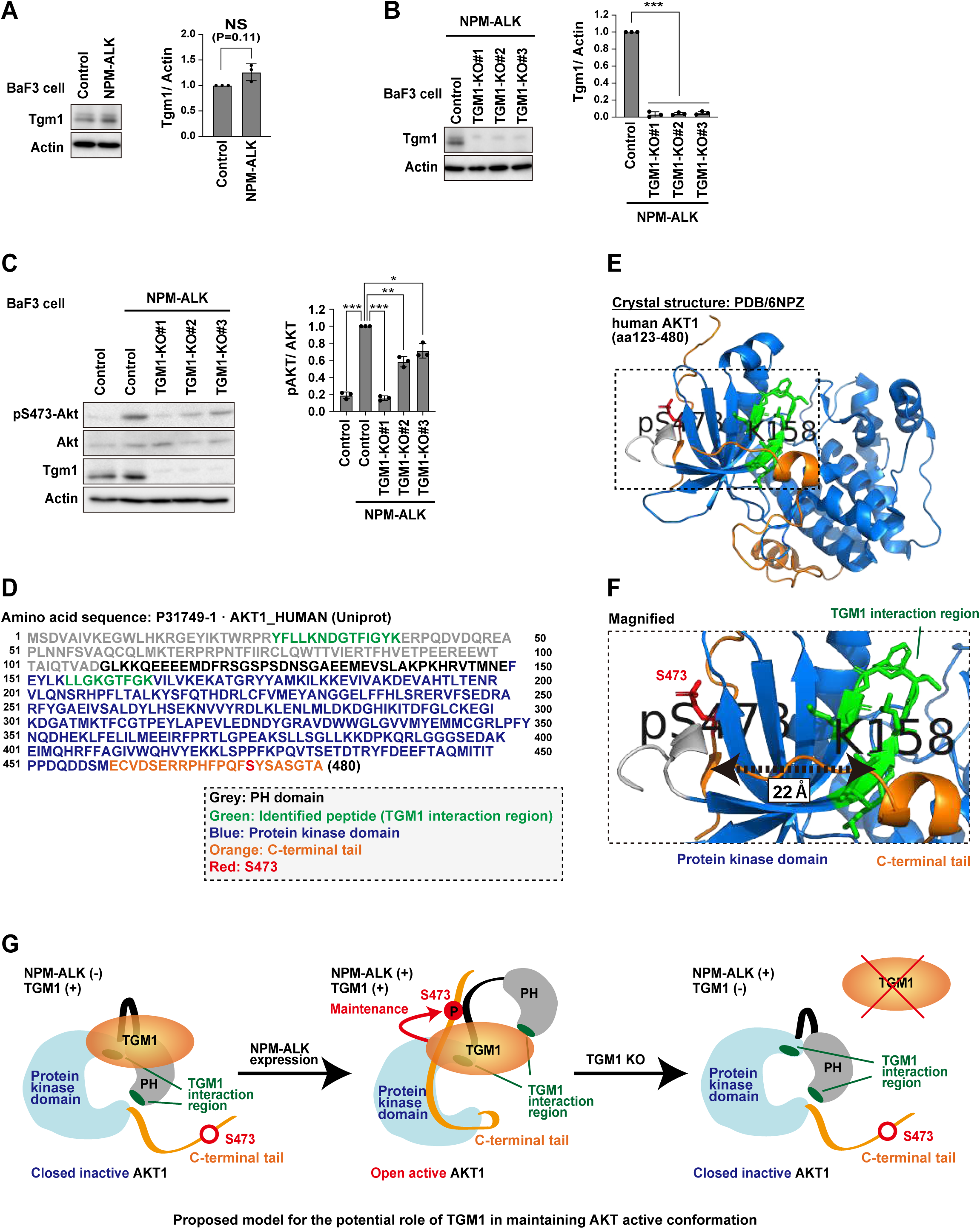
Association of TGM1 with AKT signaling in NPM-ALK–expressing cells. (A) Western blot analysis of TGM1 expression in Ba/F3 cells in the presence or absence of NPM-ALK. Quantification of TGM1 levels normalized to actin is shown. (B) Western blot analysis confirming efficient depletion of TGM1 in multiple independent TGM1 knockout clones. Quantification of TGM1 levels normalized to actin is shown. (C) Western blot analysis of AKT phosphorylation (Ser473) and total AKT levels in control and TGM1-deficient Ba/F3 cells expressing NPM-ALK. Quantification of pAKT normalized to total AKT is shown. (D) Schematic representation of the human AKT1 amino acid sequence (UniProt: P31749-1). The peptide region identified by proximity labeling analysis and associated with TGM1 interaction is highlighted in green. The protein kinase domain is shown in blue, whereas the C-terminal tail region is indicated in orange. Ser473 is highlighted in red. (E) Structural model of human AKT1 generated using PyMOL based on a published AKT1 crystal structure (PDB: 6NPZ). The identified peptide region (green), kinase domain (blue), C-terminal tail (orange), and Ser473 residue (red) are shown. (F) Magnified view of the boxed region in panel E, illustrating the relative spatial positioning between the identified peptide region and the pS473-containing C-terminal region (∼22 Å). (G) Schematic model illustrating a potential role of TGM1 in maintaining the open active conformation of AKT1.

To further examine the relationship between the TGM1-associated AKT1 peptides and AKT regulatory regions, we mapped the peptides identified by proximity labeling onto the human AKT1 amino acid sequence (Fig. 5D). Two AKT1-derived peptides were identified in the BioID analysis. Among them, the N-terminal peptide whose association with TGM1 was enhanced by NPM-ALK was located within the N-terminal region of AKT1, whereas another peptide containing a biotinylated lysine residue localized within the kinase domain (Fig. 5D). Ser473 was positioned within the C-terminal regulatory tail of AKT1 (Fig. 5D). Because the available AKT1 crystal structure used for visualization contains the kinase domain and C-terminal regulatory region but does not include the N-terminal peptide region identified in our BioID analysis [42], we used this structure to visualize the spatial organization of the kinase domain and the Ser473-containing C-terminal tail (Fig. 5E, F). Based on these biochemical and structural observations, we additionally generated a schematic model illustrating a potential role of TGM1 in maintenance of the open active conformation of AKT1 (Fig. 5G). This model is consistent with previous structural studies proposing that C-terminal phosphorylation contributes to stabilization of the active AKT1 conformation through intramolecular regulatory interactions [42, 43].

### TGM1 deficiency induces widespread transcriptional changes in NPM-ALK– expressing cells

Because oncogenic NPM-ALK is known to drive extensive transcriptional reprogramming in malignant cells [4, 44, 45], we next investigated whether TGM1 contributes to downstream gene expression changes associated with NPM-ALK signaling. Transcriptome analysis in TGM1-deficient cells revealed widespread alterations in gene expression, with a large number of genes downregulated upon TGM1 loss, along with a smaller subset of upregulated genes (Fig. 6A, B). Quantitative assessment across independent knockout clones confirmed that the number of downregulated genes consistently exceeded that of upregulated genes, indicating a predominant effect of TGM1 loss on transcriptional activation-associated programs (Fig. 6C). These results indicate that TGM1 contributes to global regulation of gene expression in NPM-ALK– expressing cells, linking interactome remodeling to downstream transcriptional changes.

**Figure 6.**
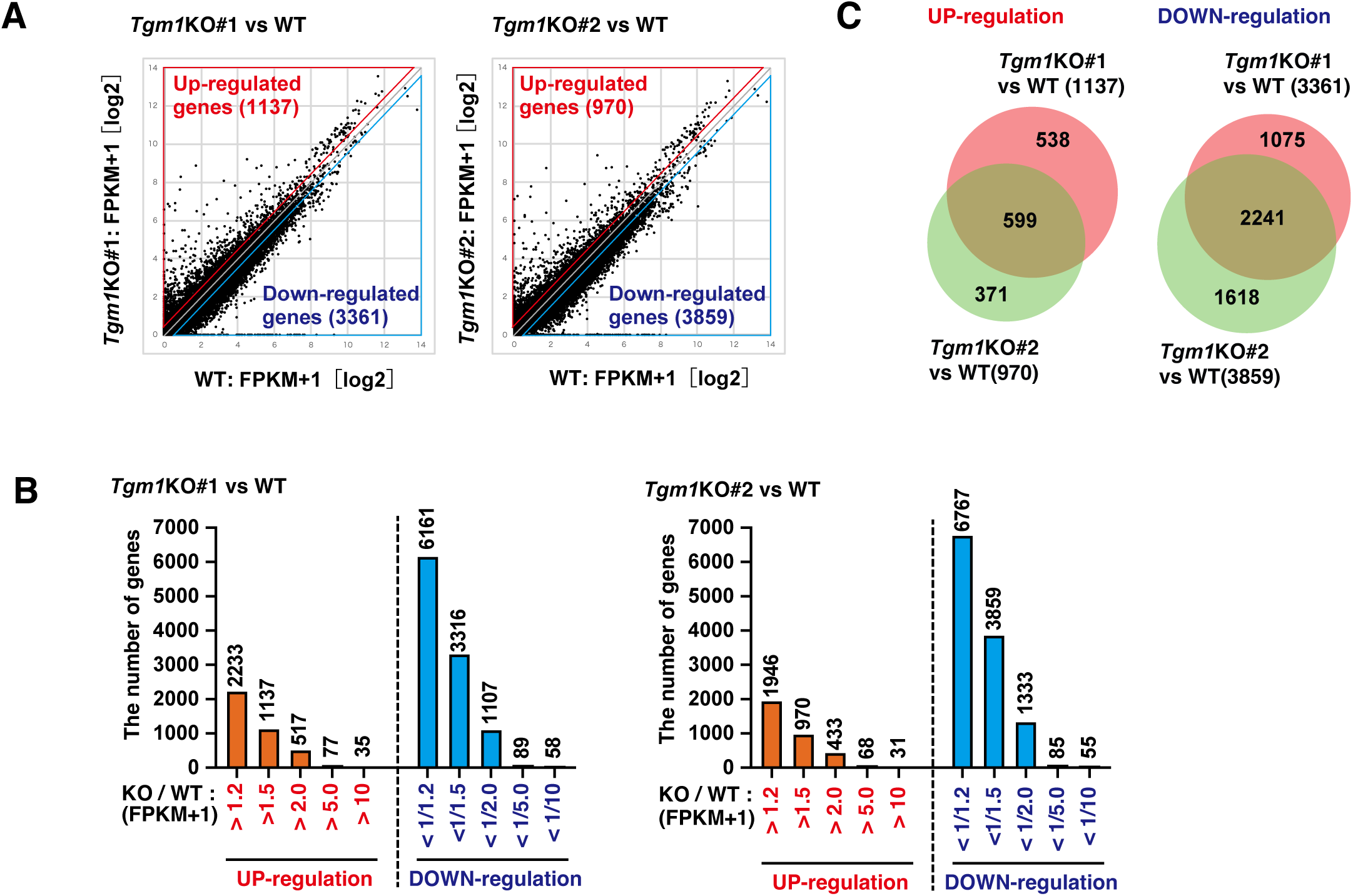
Impact of TGM1 depletion on gene expression in NPM-ALK–expressing cells. **(A)** Scatter plots showing gene expression changes in TGM1 knockout (KO#1 and KO#2) versus wild-type (WT) Ba/F3 cells expressing NPM-ALK. Each dot represents a gene. Genes were classified as upregulated or downregulated based on fold-change thresholds of >1.5 or <1/1.5, respectively. **(B)** Number of genes showing differential expression at the indicated fold-change thresholds in TGM1 knockout cells compared with WT. **(C)** Overlap of upregulated and downregulated genes between independent TGM1 knockout clones.

### TGM1 regulates transcriptional programs associated with oncogenic phenotypes

To further define the biological significance of TGM1-dependent transcriptional changes, we performed pathway enrichment analysis of differentially expressed genes in TGM1-deficient cells. Gene ontology analysis revealed that genes upregulated upon TGM1 loss were significantly enriched in apoptotic processes and stress response pathways, whereas downregulated genes were strongly associated with cell cycle progression and mitotic processes (Fig. 7A, B). Consistent with these findings, hallmark gene set analysis demonstrated enrichment of p53 signaling and apoptosis-related pathways, along with a reduction in MYC target gene expression in TGM1-deficient cells (Fig. 7C). Furthermore, representative genes involved in stress responses, such as Gadd45a and Ddit3, were upregulated, whereas key regulators of proliferation, including Mcm4 and Plk1, were downregulated (Fig. 7D). These results suggest that TGM1 contributes to the regulation of transcriptional programs that are associated with proliferation and survival while limiting stress and apoptotic responses in NPM-ALK–expressing cells.

**Figure 7.**
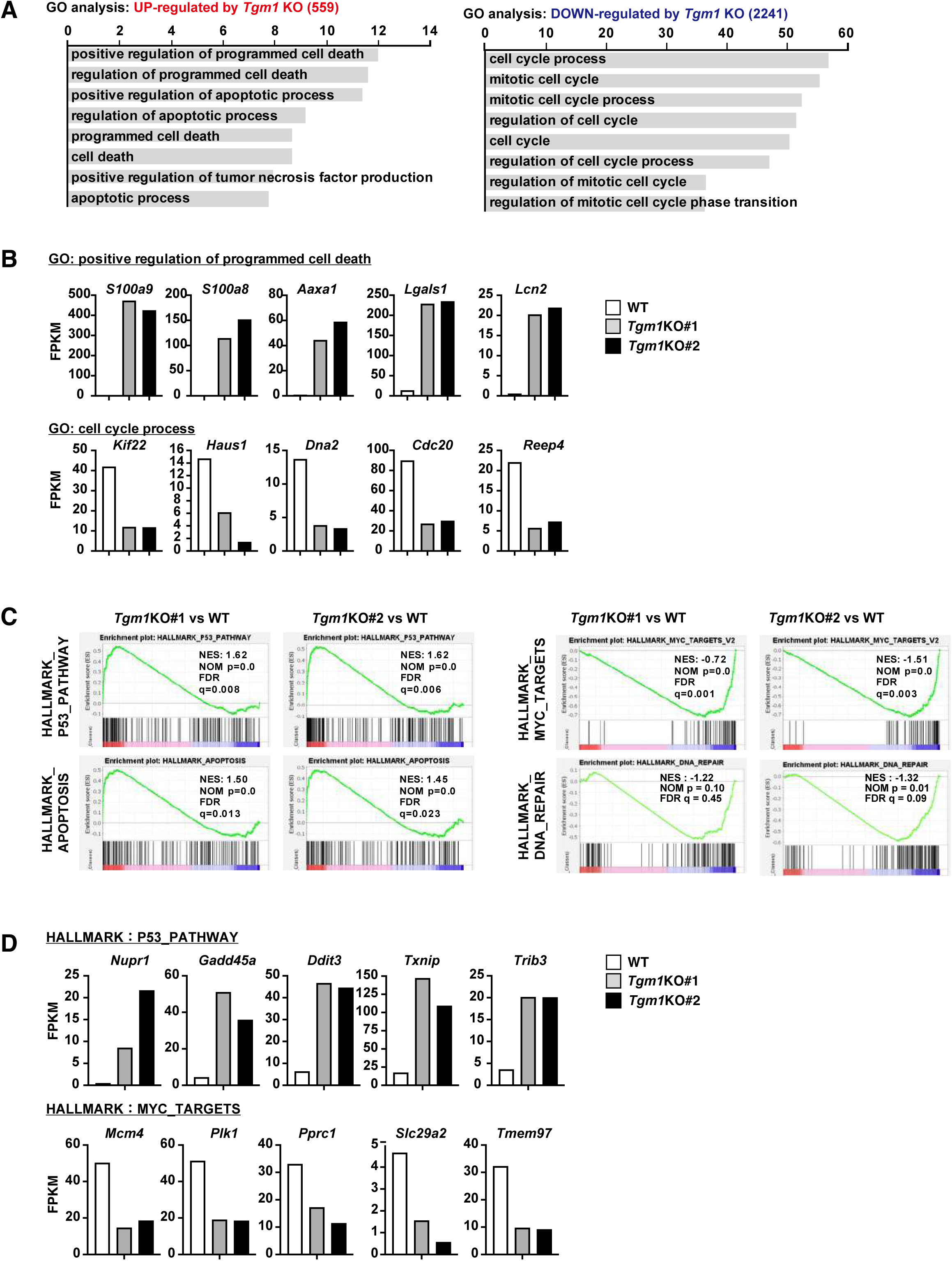
Transcriptional programs altered by TGM1 depletion in NPM-ALK– expressing cells. (A) Gene ontology (GO) analysis of genes upregulated or downregulated in TGM1 knockout cells compared with wild-type cells. (B) Expression levels of representative genes associated with programmed cell death and cell cycle pathways in TGM1 knockout and wild-type cells. (C) Gene set enrichment analysis (GSEA) of hallmark pathways, including p53 signaling, apoptosis, MYC targets, and DNA repair, in TGM1 knockout versus wild-type cells. (D) Expression levels of representative genes from p53 pathway and MYC target gene sets in TGM1 knockout and wild-type cells.

### TGM1 contributes to the proliferation of NPM-ALK–expressing cells

To evaluate the functional contribution of TGM1 to NPM-ALK–driven cellular phenotypes, we assessed cell proliferation in TGM1-deficient Ba/F3 cells. Loss of TGM1 resulted in a marked reduction in cell proliferation over time compared with control cells expressing NPM-ALK (Fig. 8A–D). This proliferative defect was consistently observed across multiple independent TGM1 knockout clones, indicating that TGM1 supports proliferative capacity in this context. In contrast, cell viability was not significantly affected by TGM1 depletion (Fig. 8E), indicating that the primary effect of TGM1 loss is on proliferation rather than cell survival.

**Figure 8.**
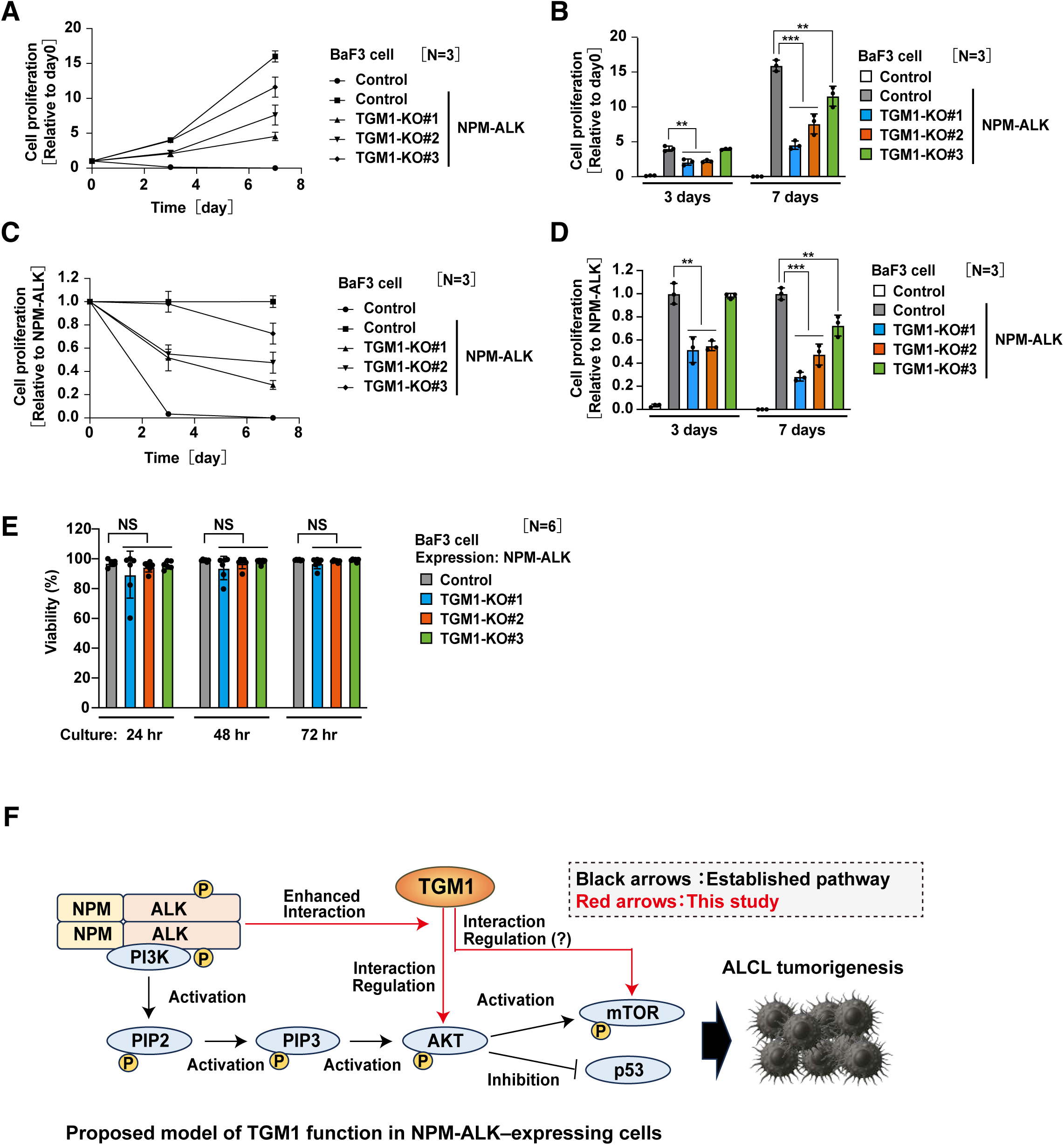
TGM1 contributes to oncogenic signaling networks required for maximal proliferative capacity. (A) Cell proliferation assay of Ba/F3 cells expressing NPM-ALK in control and TGM1 knockout clones. Cell numbers are shown relative to day 0 over the indicated time points. (B) Cell proliferation of control and TGM1 knockout cells normalized to NPM-ALK– expressing control cells. Data are presented as mean ± SD. (C) Cell proliferation assay at the indicated time points (3 and 7 days) in control and TGM1 knockout cells. (D) Quantification of cell proliferation relative to control cells at the indicated time points. Data are presented as mean ± SD. Data in panels A–D are derived from the same experiment and are presented in different formats. (E) Cell viability assay of control and TGM1 knockout cells expressing NPM-ALK at the indicated time points. Data are presented as mean ± SD. (F) Proposed model of TGM1 function in NPM-ALK– expressing cells. Established PI3K–AKT–mTOR signaling pathways are indicated by black arrows, whereas relationships involving TGM1 are shown by red arrows and represent interactions and potential regulatory effects inferred from the present study. Statistical significance is indicated as *P < 0.05, **P < 0.01, ***P < 0.001; NS, not significant.

Finally, we propose a model in which TGM1 is positioned within NPM-ALK–driven signaling networks and may contribute to interactions with components of AKT/mTOR signaling pathways (Fig. 8F). In this model, established PI3K–AKT–mTOR signaling is represented by black arrows, whereas relationships involving TGM1 are indicated by red arrows and denote interactions and potential regulatory effects inferred from the present study. Previous studies demonstrated that oncogenic NPM-ALK constitutively activates PI3K–AKT signaling and promotes downstream mTOR pathway activation, thereby supporting proliferation and survival of ALCL cells [40, 41, 46]. In particular, AKT phosphorylation and mTOR activation have been shown to play important roles in NPM-ALK–dependent oncogenic transformation and maintenance of malignant phenotypes [40, 41]. Consistent with these established signaling mechanisms, our proteomic and functional analyses identified enhanced associations of TGM1 with AKT1 and mTOR in the presence of NPM-ALK, together with reduced AKT Ser473 phosphorylation upon TGM1 depletion. Together, these findings suggest that TGM1 may contribute to signaling networks that support oncogenic cellular states.

## Discussion

In this study, we demonstrate that oncogenic NPM-ALK extensively remodels the TGM1-associated protein network, linking oncogenic signaling to downstream cellular programs (Fig. 2–4). Using proximity-dependent proteomics, we identified a broad set of TGM1-associated proteins (Fig. 2) and found that NPM-ALK expression induces a bidirectional reorganization of this network, characterized by reduced associations with proteins involved in genome maintenance (Fig. 3) and enhanced associations with components of signaling and translational pathways (Fig. 4).

These findings suggest that oncogenic signaling can reprogram protein interaction networks associated with molecules such as TGM1 (Fig. 2–4). Notably, TGM1-associated proteins were enriched in cytoskeletal and intracellular signaling pathways under basal conditions (Fig. 2), consistent with its association with intracellular signaling pathways. Upon NPM-ALK expression, this network shifts toward pathways linked to protein synthesis and PI3K–AKT signaling (Fig. 4), while associations with proteins involved in DNA repair and genome integrity are reduced (Fig. 3). This coordinated remodeling is consistent with a transition toward cellular states that favor proliferation over maintenance of genomic stability.

Proteomic and functional analyses suggested that TGM1 is linked to AKT signaling in NPM-ALK–expressing cells (Fig. 5). Consistent with the proteomic data showing increased association of AKT-related proteins within the TGM1-associated network upon NPM-ALK expression (Fig. 4), TGM1 deficiency resulted in reduced AKT phosphorylation without affecting total AKT levels (Fig. 5). These findings suggest that TGM1 may be linked to AKT signaling in this context. In addition, proximity labeling identified multiple AKT1-derived peptides associated with TGM1, including an N-terminal peptide whose association was enhanced by NPM-ALK expression and another peptide containing a biotinylated lysine residue within the kinase domain (Fig. 5D). Although these findings do not establish direct molecular interactions, they raise the possibility that TGM1-associated regions may influence structural or conformational regulation of AKT1. Consistent with this interpretation, previous structural studies demonstrated that phosphorylation of the AKT1 C-terminal tail contributes to stabilization of the active kinase conformation through intramolecular regulatory interactions [42]. Our schematic model therefore proposes a potential role for TGM1-associated interactions in maintenance of AKT activation (Fig. 5G). Previous structural studies have demonstrated that phosphorylation of the AKT1 C-terminal hydrophobic motif promotes intramolecular interactions between the C-terminal tail and the kinase domain, thereby stabilizing the active AKT conformation [42, 47]. More recent studies further showed that Ser473 phosphorylation contributes to release of PH domain-mediated autoinhibition through conformational regulation of AKT1 [43, 48]. Although our findings do not establish a direct mechanistic interaction between TGM1 and AKT1, they are consistent with a model in which TGM1-associated regions may contribute to maintenance of the active AKT conformation. However, the precise nature of this association remains to be determined. Proximity-dependent labeling captures both direct and indirect associations do not distinguish between direct binding and indirect interactions within protein complexes. Therefore, further studies will be required to clarify the molecular basis of the association between TGM1 and AKT signaling components.

Although our findings support an association between TGM1 and AKT signaling, the biological effects of TGM1 deficiency are unlikely to be explained solely by alterations in AKT phosphorylation. Given the broad remodeling of TGM1-associated protein networks observed upon NPM-ALK expression, additional signaling or regulatory pathways may also contribute to the phenotypic effects associated with TGM1 loss.

Together, our results propose a model in which oncogenic NPM-ALK reprograms TGM1-associated protein networks, thereby linking changes in protein association landscapes to downstream cellular phenotypes (Fig. 8F). By integrating proteomic and functional analyses, this study provides a framework for understanding how oncogenic signaling reshapes protein networks to coordinate signaling and transcriptional outputs.

More broadly, these findings suggest that modulation of protein-associated networks may represent an important mechanism by which oncogenic pathways control cellular states. Further studies will be required to determine whether TGM1 plays similar roles in other oncogenic contexts and to define the molecular mechanisms underlying its association with signaling pathways.

## Acknowledgments

The authors thank Kohei Nishino (Tokushima University) for his expert technical assistance with the proteomic analysis. The authors thank Junko Yamada for maintaining the shared equipment platform and for her logistical support. The authors thank Dr. Tago and Dr. Nakazawa (Division of Hygienic Chemistry, Faculty of Pharmacy, Keio University) for their general support. This work was supported in part by JSPS KAKENHI (Grant Number 23K07851); the Kobayashi Foundation; the Friends of Leukemia Research Fund (Takaku Fumimaro Award); the Japanese Society of Hematology; the Keio University Academic Development Fund; the Keio University Fukuzawa Fund; the Chemo-Sero-Therapeutic Research Institute (Kaketsuken); the Suzuken Memorial Foundation; the Mochida Memorial Foundation; and the Takeda Science Foundation; Medical Research Center Initiative for High Depth Omics. ChatGPT (OpenAI) was used under author supervision for minor revisions, consistency checking, and idea exploration. All content was verified by the authors.

## Authorship Contributions

S.T and K.A. contributed equally to this work. S.T performed most of the experiments, analyzed the data, and prepared the figures. K.H. and Y.T. assisted with experiments. H.K. conducted the mass spectrometry-based proteomic analysis. K.A. supervised the project, secured funding, and wrote the manuscript. All authors reviewed and approved the final manuscript.

